# Spatio-temporally specific transcranial magnetic stimulation to test the locus of perceptual decision making in the human brain

**DOI:** 10.1101/304063

**Authors:** Bruce Luber, David C. Jangraw, Greg Appelbaum, Austin Harrison, Susan Hilbig, Lysianne Beynel, Tristan Jones, Paul Sajda, Sarah H. Lisanby

**Author notes:** Correspondence, Susan Hilbig, Department of Psychiatry and Behavioral Science, Duke University School of Medicine, 40 Medicine Circle, Durham, NC 27705.

## Abstract

Previous research modeling EEG, fMRI and behavioral data has identified three spatially distributed brain networks that activate in temporal sequence, and are thought to enable perceptual decision-making during face-versus-car categorization. These studies have linked late activation (>300ms post stimulus onset) in the lateral occipital cortex (LOC) to object discrimination processes. We applied paired-pulse transcranial magnetic stimulation (ppTMS) to LOC at different temporal latencies with the specific prediction, based on these studies, that ppTMS beginning at 400ms after stimulus onset would slow reaction time (RT) performance. Thirteen healthy adults performed a two-alternative forced choice task selecting whether a car or face was present on each trial amidst visual noise pre-titrated to approximate 79% accuracy. ppTMS, with pulses separated by 50ms, was applied at one of five stimulus onset asynchronies: -200, 200, 400, 450, or 500ms, and a sixth no-stimulation condition. As predicted, TMS at 400ms resulted in significant slowing of RTs, providing causal evidence in support of LOC contribution to perceptual decision processing. In addition, TMS delivered at -200ms resulted in faster RTs, indicating early stimulation may result in performance enhancement. These findings build upon correlational EEG and fMRI observations and demonstrate the use of TMS in predictive validation of psychophysiological models.

## Introduction

The human brain is uniquely adept at interpreting visual input with a remarkable ability to process features, objects, and scenes, while performing complex categorizations in the blink of an eye. These extraordinary abilities are at the core of human visual cognition, and there has been a concerted effort from systems and cognitive neuroscientists to elucidate the underlying processes and neural mechanisms that enable perceptual decision making (PDM; Heekeren et al 2008). This effort has been supported, in no small part, by the powerful combination of electrophysiological and hemodynamic measures of brain function with complementary spatial and temporal sensitivity (e.g. Di Russo & Pitzalis 2013; Ales, Appelbaum, Cottereau, & Norcia, 2013)

In one particularly fruitful line of research, Philiastides, Sajda and colleagues combined electroencephalography (EEG) and functional magnetic resonance image (fMRI) collected during variants of a face/car discrimination task to characterize distributed networks that activate in sequence during PDM. In particular, through a series of three studies (Philiastides, Ratcliff, & Sajda, 2006; Philiastides & Sajda, 2006, 2007) these authors utilized single trial logistic regression on EEG, drift diffusion modeling of behavioral data, and EEG-informed fMRI analysis to ascertain the cortical origins of three temporally specific neural networks sensitive to different elements of the task parameterization.

These studies utilized a behavioral task in which participants discriminated face images from car images that were degraded in perceptual clarity through scrambling of spatial phase with values from 20% to 45% coherence. In a first study, sixty-channel EEG data recorded during performance of the task was analyzed on a single trial basis using logistic regression to maximally distinguish face and car trials (Philiastides & Sajda, 2006; Philiastides, Ratcliff, & Sajda, 2006). Psychometric functions relating performance accuracy and coherence levels were statistically indistinguishable from neurometric functions relating the strength of classification to coherence levels, suggesting the EEG reflected the workings of the neural substrate of the categorization. The best matches of these functions occurred in an early latency window, centered on 170 ms from stimulus onset, which corresponds to the N170 ERP component well-known to be involved with stimulus categorization, and a later window, beginning after 300 ms latency, which formed even better matches with the performance data than the early one. In fact, both the onset latency and the duration of the later EEG component increased with discrimination difficulty (i.e., with decreased phase coherence), relationships not found in the early component. Further, the early component was just as active when evoked during a simple red/green discrimination, while the late component was only evoked when the more difficult degraded face/car categorization was made. Using a drift diffusion model to link the accumulation of information over time to decision choices, it was found that the estimated drift rate in the model was strongly correlated with the strength of discrimination estimated from the EEG data of the late, but not the early, component. Furthermore, a third component, peaking around 220 ms, was also identified and whose activity was found to be closely bound to the activity of the late component and to stimulus difficulty: its amplitude was inversely proportional both to the stimulus evidence in the model and to the onset of the late component. Overall, the evidence from these studies indicated three components of neural activity: an early one involved in initial perceptual processing, and two later ones that closely linked to PDM.

A third study from this group utilized fMRI to ascertain the cortical origins of each of the three temporally specific EEG components identified in the first two studies (Philiastides & Sajda, 2007). Using the previous EEG results as fMRI regressors, they identified a spatio-temporal cascade of activity in three spatially-distributed networks, with contributions from the fusiform face area and superior temporal sulcus associated with the early component, a network of mainly frontal attention-related areas mediating the difficulty-dependent second component, and the involvement of the lateral occipital cortex (LOC) with the late component. This fMRI study therefore tied together findings from the other two studies to link the spatial and temporal patterns of activity in networks underlying decision-making in uncertain conditions by correlating behavioral performance with network activity.

While these studies provide strong evidence towards the involvement of discrete brain networks in different stages of PDM, their findings are correlational, and do not provide definitive evidence of causal brain-behavior relationships. In contrast to EEG and fMRI, transcranial magnetic stimulation (TMS) is a type of non-invasive brain stimulation that can be used to establish such causal links, given its ability to selectively perturb neural information processing and measure the effects on behavior. In particular, the exacting psychophysical, electrophysiological and imaging work of Philiastides, Sajda and colleagues lends itself to a very specific test of their dynamic neurophysiological model. Namely, that a pair of disruptive TMS pulses, applied during a time window beginning at 400ms after the stimulus onset would inject neural noise (as developed in Kammer et al., 1998) during the sensitive period related to the face/car PDM and would be expected to slow down discrimination processing, resulting in a longer reaction time. The TMS timing parameters used in our prediction were carefully based on the findings of Philiastides, Sajda and colleagues. First, by previously titrating phase coherence to a specific accuracy level of 79% for both face and car stimuli using an adaptive staircase, a phase coherence level could be applied in each subject that would be expected to push the onset latency of the late component to an approximate time of 400 ms, based on the temporal relationship of phase coherence with late component onset latency found in Philiastades and Sajda (2006) and Philiastades et al. (2006). Second, a pair of TMS pulses, separated by 50 ms, was used to fit within a 60 ms time window, which was found to be the window duration that best discriminated the late component in the single trial EEG analyses of Philiastides et al. (2006). Third, based on the relationship of EEG activation and drift rate in the diffusion model of behavioral data during the late component found in Philiastides et al. (2006), we expected the random neural excitement added by TMS in the 400 ms window would approximate the addition of noise to the diffusion model, resulting in slower processing and thus a lengthening of reaction times.

In accord with this prediction, participants in the present study performed a speeded version of the face/car task with paired pulse TMS (ppTMS) applied with a 50 ms interstimulus interval introduced at a number of latencies spanning from 200 ms before stimulus presentation, to 450 ms post stimulus. ppTMS was applied to the LOC, the major source of the late component activity in the Philiastides and Sajda MRI study, and it was hypothesized that this stimulation during the active phase of perceptual discrimination would result in impaired performance relative to stimulation at 500 ms, a latency the previous work indicated was past the completion of the late component. As such, the current design utilizes a chronometric approach to perturb neural function across the range of times specifically identified as important for PDM, providing a causal test of these past correlational relationships.

## Materials and Methods

### Participants

Fifteen healthy volunteers were recruited and provided written informed consent for the study, which was approved by the Institutional Review Board of the Duke University Medical Center. Two participants dropped out for scheduling reasons, leaving 13 completing the full study. These 13 individuals had a mean age of 24.6 ± 2.8 years and consisted of five female and eight males. Participants had normal, or corrected-to-normal, vision and were native English speakers. Potential participants were excluded if they had current or past Axis I psychiatric disorder including substance abuse/dependences, as determined by the MINI International Neuropsychiatric Interview, English Version 5.0.0 DSM-IV (Sheehan D, 2006), or neurological disease as determined by the TMS Adult Safety Screen (Keel, Smith, & Wasserman, 2001). All participants were screened for substance abuse with urine drug screens and women of childbearing capacity were screened with urine pregnancy tests. Following screening, participating subjects were introduced to the task then returned for three more visits within the next 2 weeks; an MRI session, and two TMS sessions utilizing two counterbalanced targets.

### Stimuli

A set of 12 face (chosen from the Max Planck Institute face database, http://faces.kyb.tuebingen.mpg.de) and 12 car images were used, with both sets having equal numbers of front and side views. All images were rendered in grayscale with 8 bits/pixel, were 512 × 512 pixels in size, and were equated for spatial frequency, luminance, and contrast (for a more complete description, see Philiastides and Sajda, 2005). Stimuli were presented on a Dell UltraSharp 20.1” monitor at a distance of 100 cm, using E-Prime software (Psychological Software Tools, Pittsburgh, PA). The phase spectra of the face and car images were manipulated to generate sets of stimuli with varying degrees of phase coherence (**Figure 1A**).

**Figure 1:**
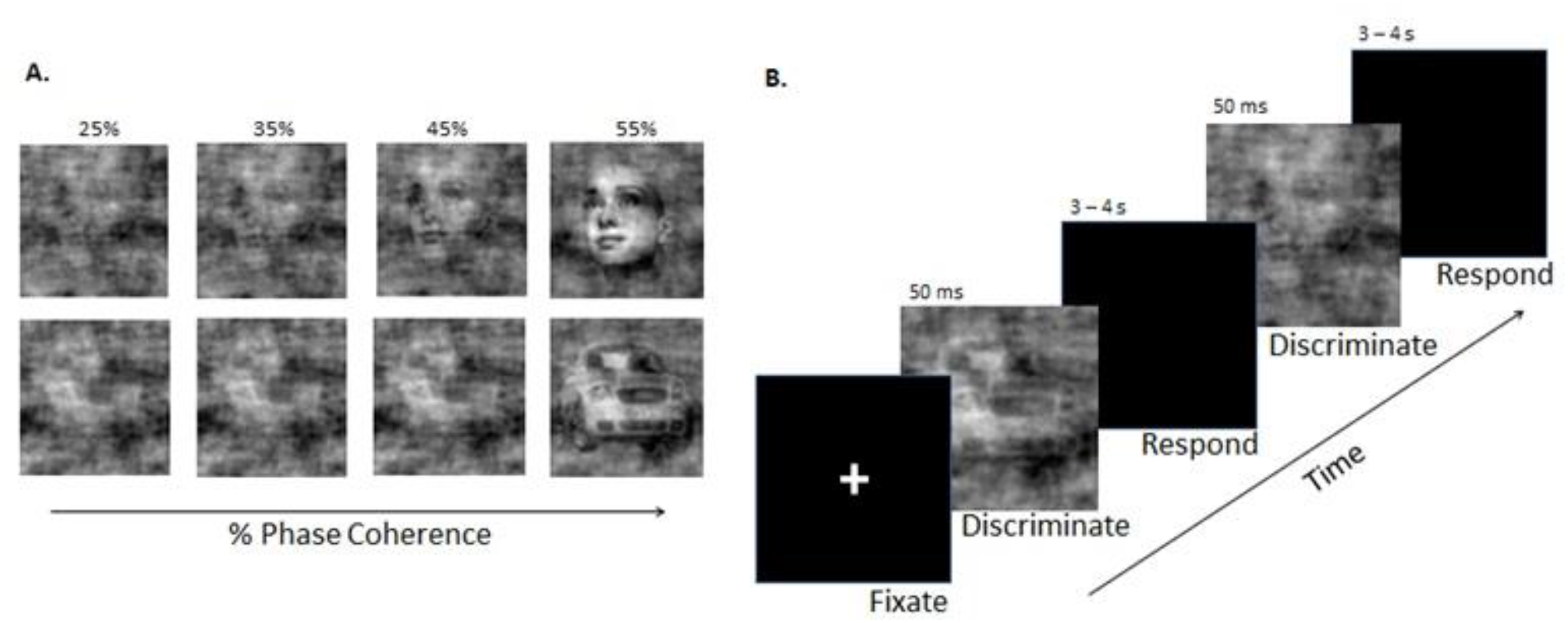
A) Examples of face and car stimuli with various levels of noise added by manipulation of the phase coherence in the image. B) Schematic illustration of the Visual Discrimination Task. Phase coherence varied across trials according to a staircase schedule and determined by each individual’s performance.

### Visual Discrimination Task

Participants were seated, resting forward on a cushioned massage chair facing the monitor with his/her head resting in the open headrest of the chair. Using a Cedrus RB-830 Response Pad (San Pedro, CA), participants were instructed to respond, as quickly and as accurately as possible, whether a face or car image was displayed on the screen. The order of presentation for the car and face stimuli was randomized across trials. Stimuli were presented for 50 ms, followed by a blank screen over an inter-trial interval that was randomly selected with a uniform distribution between 3000 and 4000 ms, during which the participant had to respond (**Figure 1B**).

### Psychometric Staircase

At the beginning of each session, participants performed the visual discrimination task until their performance accuracy became stable at 79% for both stimulus types. This served two purposes: to reduce variability between individuals in task performance, and, more importantly, to push subjects to perform in a more difficult range of coherence levels that was previously shown to correspond to an onset of the late component of approximately 400 ms (Philiastides and Sajda, 2006; Philiastides et al., 2006).

This was achieved using a staircasing algorithm (Levitt, 1970), which dynamically changed the image coherence in successive stimuli, independently for faces and cars. To that end, two interleaved staircase functions were run across a block of trials: one containing only faces and the other containing only cars, with trials from the two staircases interspersed randomly. The initial stimuli in both staircases were presented at the easiest (i.e., most discriminable) level used, with 50% phase coherence for both faces and cars. Participants were asked to respond as quickly as possible via button presses whether the image was a face or car. If the participant answered three consecutive trials from one staircase correctly, the difficulty of that staircase increased (i.e., decreased coherence in the next stimulus), while a single incorrect response decreased the difficulty level one step. This “three-up, one-down” procedure theoretically converges at a participant performance level of 2^^(-1/3)^, or approximately 79.4% correct. Each time the change in difficulty changed sign it was considered a “reversal.” Before the 4th reversal, the difficulty changed in increments of 10% phase coherence. Between the 4th and 10th reversals, the difficulty changed in increments of 3%. After the 10th reversal, the difficulty changed in increments of 1%. The staircase paradigm also served as additional practice for the participants, allowing them to reach constant performance before the final difficulty was calculated. For this reason, the paradigm ended after both staircases had completed 25 reversals. For each staircase, the coherence levels over the last 10 reversals were averaged to estimate threshold performance for this perceptual discrimination task. Participants were allowed to take a break every 5 minutes for as long as needed, after which they resumed the task at their previous performance level.

### MRI acquisition and analysis

Prior to the TMS sessions, MRIs were obtained for use in later TMS targeting. MRI scanning was conducted on a 3T General Electric scanner at Duke University (Durham, NC) using an 8HRBrain coil configuration and spin echo pulse sequence. Whole-brain T1-weighted MRI scans were collected and submitted to automated tissue segmentation and definition of anatomical regions of interest (ROIs). The anatomical MRI was acquired using a 3D T1-weighted echo-planar sequence (TR = 8.208 ms, TE = 3.22 ms, FOV = 240*240, slice thickness = 1.6mm). During this scan, the participants viewed a blank screen and relaxed. Anatomical images were skull-stripped and reoriented from LPS (Left, Posterior, Superior) to LAS (Left, Anterior, Superior) orientation using FMRIB Software Library (FSL v.5.0). The left and right LOC coordinates were first defined on the Montreal and Neurological Institute Template according to group fMRI analyses using the face/car discrimination task (Philiastades and Sajda, 2007). These coordinates (left LOC: -42, -88, -10, right LOC: 46, -86, -8) were then registered on each individual brain using FSL affine linear regression tool. Final coordinates were uploaded on a frameless stereotaxic neuronavigation system (Brainsight, Rogue Research, Canada) to guide the TMS coil positioning.

### TMS sessions

Two TMS sessions were run on separate days, each lasting approximately 2 hours. Each session consisted of the staircasing procedure, described above, to obtain coherence levels for each image type on that day. This staircasing procedure was followed by 6 blocks of visual discrimination task, with TMS applied to either left or right LOC. The other LOC site was targeted in the second session, with the site order counterbalanced across participants. While previous studies have found both left and right LOC to be active during visual discrimination tasks (Ales, Appelbaum, Cottereau, & Norcia, 2013; Philiastides & Sajda, 2007), left and right sites were stimulated separately here in order to evaluate laterality effects.

TMS was delivered using a Cool-B65 Butterfly figure-8 coil powered by a MagPro X100 stimulator (MagVenture, Farum, Denmark), and positioned using the neuronavigation system and SmartMove robotized TMS coil positioning system (ANT, Enschede, Netherlands) allowing 300 ms recovery and 1-3 millimeter precision. Stimulus intensity was set at 100% of the participant’s resting motor threshold, collected at the beginning of the first TMS session and defined as the minimum magnetic intensity needed to evoke motor potentials of at least 50μV recorded via EMG from the first dorsal interosseus muscle (FDI) of the participant’s right hand in at least 5 out of 10 stimulations (Rossi, Hallett, Rossini, Pascual-Leone, & Safety of TMS Consensus Group, 2009).

One-sixth of the task trials in a block of trials were no-TMS trials in which no TMS stimulation was delivered, and the other five-sixths were TMS trials. In each TMS trial, paired TMS pulses separated by 50 ms were delivered. Past research has indicated ppTMS can be used to disrupt visual processing in LOC (Ellison & Cowey, 2007; Mullin & Steeves, 2011; Pitcher, Garrido, Walsh, & Duchaine, 2008) with an acute disruptive effects lasting about 50 ms (e.g., Amassian et al., 1989). The 50 ms timing of the pulses in the present study were chosen to disrupt processing over a similar time range as the 60 ms time window in single trial EEG analyses that was found to be the best window duration to capture the activation and psychophysiological relationships of the late component (Philiastides et al., 2006).

During the paired pulse stimulation, the first of the two pulses was time-locked to the onset of the visual stimulus at one of five stimulus onset asynchronies (SOAs): -200, 200, 400, 450 or 500 ms. Paired-pulse TMS applied at 500 ms SOA was defined as the control condition, given that the previous research indicated that the cognitive processing associated with the late component at LOC is expected to be completed at this time (Philiastides and Sajda, 2006; Philiastides et al., 2006). Such a temporal control condition has been used in the field since its inception (e.g., Amassian et al., 1989), and by our own group in previous visual studies (Matthews et al., 2001; Luber et al., 2007). This control condition presents several advantages compared to using an active control site or a passive sham condition: (1) the different SOAs can be randomly interspersed within the same block of trials, with the subject having little awareness of their difference and no ability to predict them; (2) these conditions feel the same to the participant; and (3) the different conditions stimulate the same nervous tissue. Moreover, a second control site is unnecessary in the present case, as we are specifically testing the prediction that TMS to LOC at a specific time (relative to another) will slow RT.

The choice of SOA (or no-TMS) in a given trial was made pseudo-randomly, constrained so that the total number of each of the six conditions was equal in each block of trials. There were 162 trials in each of the six blocks, and thus 162 trials for each of the six conditions.

### Analysis

Median RTs in correct trials and accuracy were calculated for each stimulus type at each SOA. A 2 × 2 × 6 repeated measures MANOVA was performed for both these two measures, with factors of Site (left, right), Stimulus type (face, car), and SOA (-200, 200, 400, 450, 500, no-TMS). Post hoc analyses were performed for RT and accuracy measures to test the prediction that TMS at the 400 and 450 ms SOAs (Bonferroni corrected) had a deleterious effect on performance relative to the 500 ms

SOA, a latency at which the processing was expected to be completed. Exploratory post hoc tests were done between the 500 ms condition and 1) the no-TMS condition, to provide evidence that TMS at 500 ms had no effect on performance; 2) the 200 ms condition, to observe any performance effects at the latency of the second component, which was related to difficulty and expected to be active at the stimulus coherence levels employed (Philiastides et al., 2006); and 3) the -200 ms condition, to observe whether a pulse prior to the onset on the trial affected performance, as has occurred in other TMS studies (e.g., Grosbras and Paus, 2003). Effect sizes were estimated using Cohen’s d, calculated in a repeated-measures situation as the t-value obtained, divided by the square root of the degrees of freedom (Rosenthal and Rosnow, 1991).

## Results

### Coherence Thresholds

The titrated coherence thresholds for face and car stimuli were stable across the two sessions and were similar for both types of stimuli. The group mean coherence for car stimuli (34.0%, +/- 6.3) was higher than for faces (29.5% +/- 5.6), but not significantly so. A 2×2 repeated measures ANOVA on the threshold estimates, looking across the two sessions and the two stimulus types (face, car) showed no significant main effect for either factor. There was a significant Session by Stimulus Type interaction (F(1,12) = 7.0, p < 0.02), due to a decrease in average coherence needed for car stimuli between the first and second sessions, although a post hoc paired t-test was not significant for this difference. The titration procedure proved successful, in that the coherence levels used for each participant produced overall accuracy levels close to the expected staircase accuracy during the experimental sessions (group mean and SD in the no-TMS condition 75.6% +/- 7.9) with no difference between face (75.6% ± 14.6) and car stimuli (75.6% ± 12.2).

### No-TMS vs SOA 500 Conditions

To test the reliability of the TMS applied 500 ms after the stimulus onset as a control condition, we compared performance obtained in this condition to performance in the no-TMS condition where stimuli were presented without TMS stimulation. Our results did not reveal any differences between these two conditions on either accuracy (t(12) = 1.9, p=0.29) or on reaction times (t(12)= 1.2, p=0.26). This lack of difference provides validation for the use of the 500 ms SOA as a control condition, as expected from the EEG data of Philiastades & Sajda (2006) and Philiastades et al. (2007), in which activation related to the late component was complete by 500 ms.

### Reaction Time

Behavioral analyses revealed a significant effect of SOA (F(5,8) = 16.5, p < 0.0005), but no main effects of Site or Stimulus Type, and no significant interactions (**Figure 2**). Bonferroni corrected analyses showed that when TMS was applied at 400 ms SOA, RT are slower relative to TMS applied at 500 ms SOA baseline (t_12_ = 2.9, p < 0.015; Cohen’s d = 0.84). However, when TMS is applied at -200 ms SOA, RT are significantly faster compare to the 500 ms SOA (t_12_ = 4.6, p < 0.0005; Cohen’s d = 1.33).

**Figure 2.**
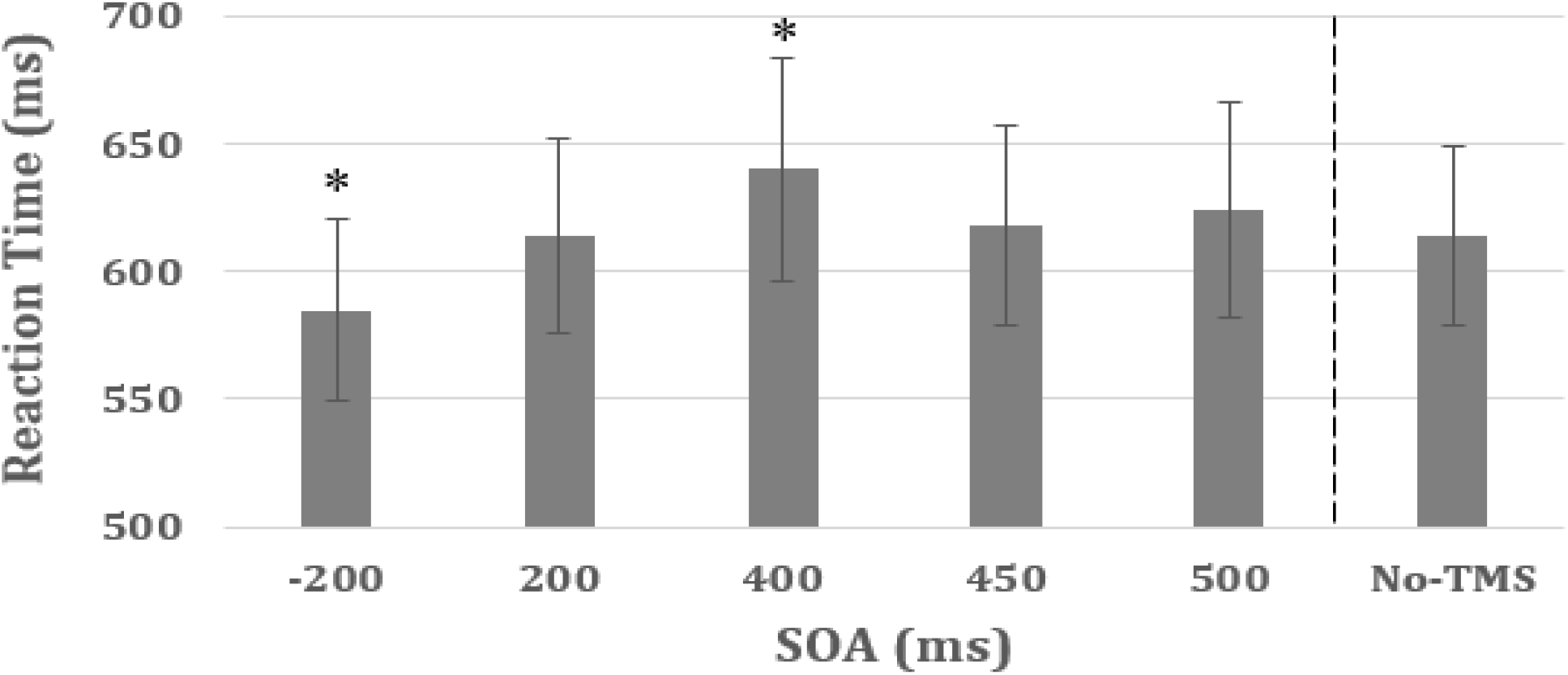
Median reaction time for correct trials, averaged across the left and right LOC stimulation sites and the face and car stimulus types. Error bars represent standard errors.

### Accuracy

The analysis revealed no significant main effects of Site or Stimulus Type (**Figure 3**). There was, however, a significant main effect of SOA (F(5,8)=3.2, p < 0.015) and a significant interaction between Stimulus Type and SOA (F(5,8) = 7.4, p < 0.01). Post hoc t-tests indicated a decrease in accuracy at 200 ms, compared to the 500 ms SOA (t_12_ = 2.6, p < 0.012; Cohen’s d = 0.75).

**Figure 3.**
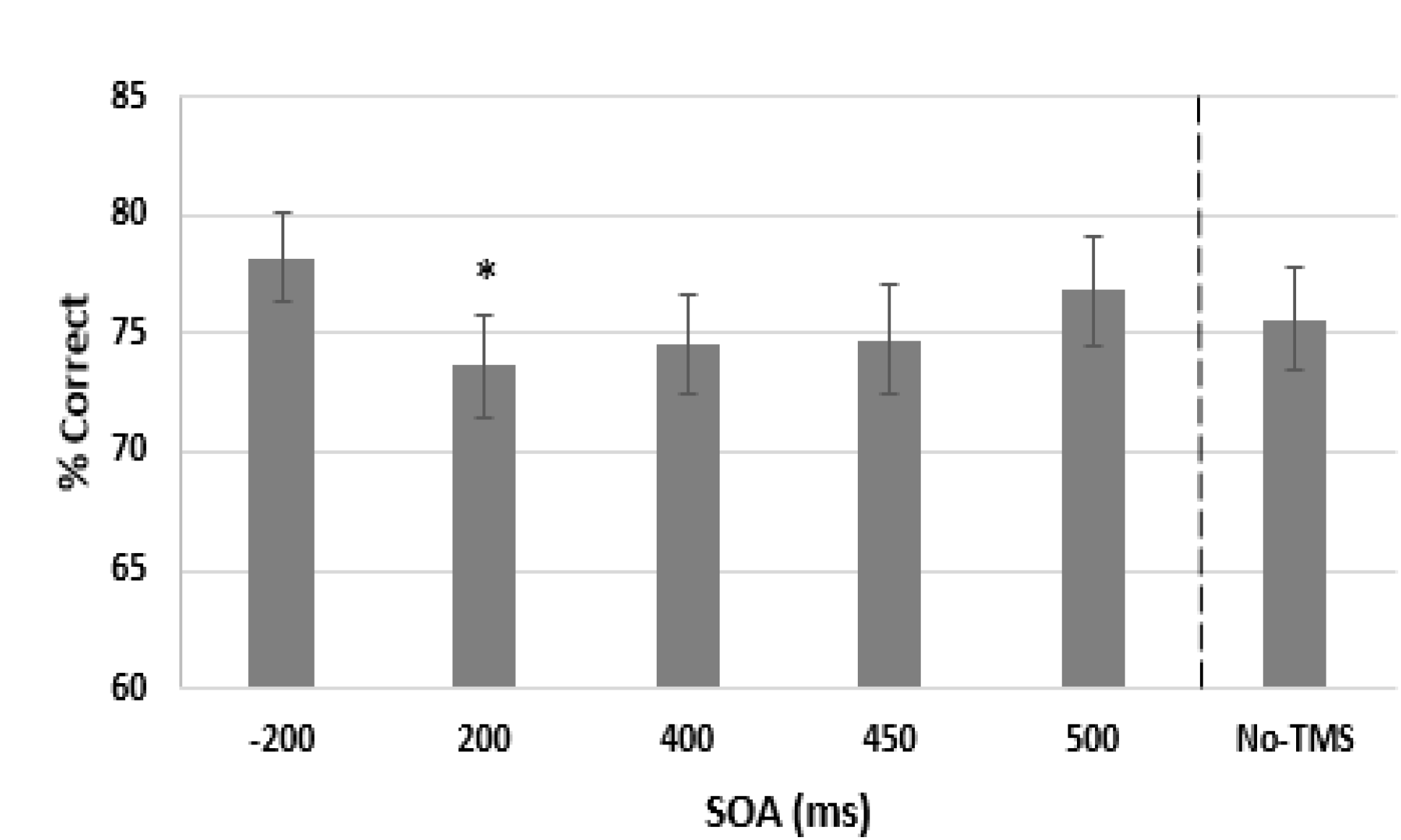
Accuracy across the SOA conditions, collapsed across the left and right stimulation sites. Error bars represent standard errors.

## Discussion

In this study, we tested the spatio-temporal model of the neural substrate of visual object decisions identified by Philiastides and Sajda. This model predicted that, at a certain level of perceptual difficulty in discriminating cars from faces (which could be controlled on an individual basis with a titration procedure), paired-pulse TMS (ppTMS) to LOC in a predicted time window (i.e., 400-450 ms latency) would interfere with decision-making processes occurring there in that window, slowing reaction time. The results did indeed reveal a significant general slowing of RT at the location and latency when the Philiastades and Sajda network model expected visual object discrimination to be occurring. Such a finding provides evidence in support of the model, and specifically that LOC plays a role in the decision process involved in object identification and PDM. Furthermore, we did not find any differences in TMS effects with either right or left LOC stimulation which also verified the relatively equal, bilateral LOC activation defined by Philiastides and Sajda model (Philiastides and Sajda, 2007).

This result is consistent with other studies showing that ppTMS applied to LOC can disrupt visual processing (Ellison & Cowey, 2007; Mullin & Steeves, 2011; Pitcher et al., 2008; Pitcher, Goldhaber, Duchaine, Walsh, & Kanwisher, 2012; Pitcher, Walsh, Yovel, & Duchaine, 2007). For example, using earlier range of SOAs than the current study, Mullin and Steeves (2011) had subjects discriminate natural versus man-made objects and scenes, while stimulating with pairs of pulses spaced 100 ms apart at SOAs of 0, 40, 80, 120, 180 and 220 ms. They found significant drops in accuracy (without affecting RT) for TMS to LOC, compared to vertex stimulation or no-TMS. These effects occurred only at early latencies: 0, 40 and 80 ms with left hemisphere stimulation and 40 ms on the right. Similarly, a second study Ellison and Cowey (Ellison & Cowey, 2007) applied ppTMS spaced 50 ms apart over LOC, at SOAs ranging from 0-350 ms, during a distance discrimination task in which subjects had to choose which of two green squares were closer in proximity to a central green square at fixation. A detrimental effect on performance was also found when TMS was applied at early SOAs of 0, 50, 150ms with a slowdown of RT compare to sham. These results suggest that LOC is involved with the early visual processing associated with the first network in Philiastades and Sajda’s model, although a simpler explanation may be that TMS to LOC at earlier latencies could disrupt processing trans-synaptically via feedback connections in the regions of the model’s first network which are active at this time.

Besides, when TMS was applied to LOC at 200 and 350ms after the stimulus onset, Ellison and Cowey (2007) found an increase in RT without affecting accuracy, although they were using a spatial discrimination task, rather than object discrimination. Interestingly, our current study also found a drop in performance when TMS was applied 200 ms after the stimulus onset, as reflected by a significant decrease in accuracy in the 200 ms SOA compared to the 500 ms SOA. The main difference between these two studies and Mullin and Steeves study is the task difficulty. In Mullin and Steeves (2011), the task involved a very easy visual discrimination, with accuracy rates without TMS of 95%, while the discrimination accuracy used in the present study and in Ellison and Cowey study (Ellison & Cowey, 2007) was titrated on an individual basis to be about 75% and 80% correct, respectively. According to Philiastades and Sajda (2007), when discrimination is difficult, a second network exerting top-down influence is activated in the 180-280 ms period. As a consequence, TMS applied around this period might have interfered with the workings of this second network.

Finally, an unexpected improvement in reaction time was found when paired-pulse TMS was applied 200 ms before the onset of the trial, as reflected by the improvement in the reaction time. It was anticipated that TMS prior to stimulus delivery would provide a second control time point, since processing of the stimulus could not yet have begun. Despite this expectation, the findings that TMS at - 200 ms SOA facilitated performance are consistent with findings of TMS performance enhancements in general (Luber & Lisanby, 2000). For example, Grosbras and Paus (2002; 2003) found that stimulation delivered in the 100 ms before the onset of a small target light increased its detectability. They suggested the TMS potentiates local neural activity for a brief period, noting that in animal studies direct electrical stimulation of neurons in the homologous visual area immediately preceding a target improved performance as well. On the other hand, because TMS also produces superficial effects including a clicking sound and mechanical vibrations passed from the coil to the scalp, intersensory facilitation may have led perceptual priming (Tereo et al., 1997). This facilitation - a speeding of RT- has been shown to occur with TMS pulses applied in the 150 ms period prior to stimulus onset. In light of these different contributions, and the possibility that TMS may differentially affect behavior when applied to different brain states, further investigation is necessary to determine whether TMS can cause enhancement of visual discrimination processing beyond possible facilitation effects and how these differ from classical perceptual priming.

The present study extends the work done by others in examining the effects of TMS on object processing in LOC, by looking for effects in a later latency range than used previously, by using more difficult-to-discriminate object stimuli that required extended PDM processing, and most importantly by specifically testing a well-developed spatio-temporal model of the neural networks involved. TMS was used to perturb neural processing at a location and latency that was expected to slow PDM processing according to a model generated by Philiastides and Sajda. The behavioral data from the face/car discrimination task had been modeled by a diffusion (or random walk) process (Ratcliff & Rouder, 1998), and fit to psychophysiological data measured at latencies between 300 and 500 ms after the visual stimulus is presented (Philiastades et al., 2006). Ratcliff’s random walk model is driven by the accumulation of information, and the cortical interference produced by TMS was expected to disrupt that process during difficult discrimination in a time period around 400 ms, slowing it down by decreasing the drift rate (size of the “steps” taken in the walk). Indeed, the slowest RTs were observed in conditions with paired-pulse TMS starting 400 ms after the visual stimulus onset, corroborating the hypothesis that the LOC is involved with object decision-making in this time window and providing evidence for the specific model of Philiastades and Sajda.

On the other hand, the reduced discrimination accuracy seen at the 200 ms SOA appears to be a qualitatively different TMS effect, suggesting interference with ongoing cortical processing of a different kind. One possibility is that the TMS is interfering with the influence on LOC of the second network in the Philiastades and Sajda model, which is active in this time period. In this model, the second network is thought to provide top-down control to aid processing when the discrimination becomes difficult. Finally, the improvement when TMS was applied before the stimulus onset need to be more investigated to define if it is due to specific TMS effect, such as a potentiation of local neural activity, or non-specific TMS effect such as intersensory facilitation.

## Conclusion

This study represents a first step in using TMS to verify an established multiple-network model of visual object discrimination, which was based on psychophysical, EEG and fMRI measurements taken during a challenging discrimination task. We were able to provide causal evidence for a prediction of the model that TMS applied to LOC at 400 ms would slow response time. In addition, we were able to observe other significant effects caused by TMS, notably a potential performance enhancement, that could be interpreted by the model and which lead to future experiments using TMS to engage the three networks posited by the model and explore their interactions. The study had a number of limitations, first that the sample size was small, leading to a need to replicate the findings in a larger group. TMS coil targeting was done using neuronavigation with individual structural MRIs and group-level functional coordinates of task activations, but one source of variability in targeting could be removed by using individual fMRI activations produced by the task in particular regions of interest. More recent findings regarding regions in lateral occipital cortex specialized for particular types of stimuli such as faces need to be taken into account in choosing task stimuli and specific cortical targets. SOAs at which TMS was not expected to affect processing, as well as no-TMS conditions, were used for control comparisons, but additional controls such as additional stimulation sites and sham TMS could be added that would improve interpretation. Nonetheless, we believe the results of this study demonstrate the usefulness of TMS in validating network models of cortical function, and in particular the Philiastades and Sajda model of visual discrimination. Further studies, especially using simultaneous TMS/fMRI to observe the immediate and long-range effects of TMS within the posited networks, provide exciting possibilities to extend this research.

